# *S*-Alk(en)yl-Cysteine Sulfoxides in Allium Species Are Excellent Acrolein Scavengers: Implications for Secondary Antioxidants in Plants

**DOI:** 10.1101/2025.08.18.670978

**Authors:** Ayako Hada, Chihiro Nozaki, Natsumi Tamura, Kenji Matsui, Yasumasa Matsuoka, Daisuke Shibata, Jun’ichi Mano

**Affiliations:** Graduate School of Sciences and Technology for Innovation, Yamaguchi University, Yoshida 1677-1, Yamaguchi 753-8515, Japan; Faculty of Agriculture, Yamaguchi University, Yoshida 1677-1, Yamaguchi 753-8515, Japan; Advanced Technology Institute, Yamaguchi University, Yoshida 1677-1, Yamaguchi 753-8515, Japan; Kazusa DNA Research Institute, 2-6-7 Kazusa-kamatari, Kisarazu Chiba, 292-0818 Japan; Science Research Center, Organization for Research Initiatives, Yamaguchi University, Yoshida 1677-1, Yamaguchi 753-8515, Japan; The United Graduate School of Agricultural Science, Tottori University, Tottori 680-8553 Japan

**Keywords:** acrolein, alliin, isoalliin, methiin, reactive carbonyl species

## Abstract

Reactive carbonyl species (RCS), such as acrolein (Acr), are generated through the degradation of lipid peroxides and exert cytotoxic effects. To identify natural RCS scavengers, we examined 80% ethanol extracts from 46 angiosperm species for Acr-trapping activity using an HPLC-based assay. Strong activities were observed in several taxa, including garlic, spinach, avocado, broccoli, and lotus. In garlic, the active metabolite was identified as *S*-allyl-L-cysteine sulfoxide (alliin), a characteristic *Allium* amino acid. Alliin and its *S*-(1*E*)-propenyl and *S*-methyl derivatives (isoalliin and methiin, respectively) trapped up to two Acr molecules at the amino group and exhibited higher activities than known scavengers such as carnosine and epigallocatechin gallate. These findings highlight *S*-alk(en)yl-L-cysteine sulfoxides as potent secondary antioxidants and suggest that structurally diverse RCS scavengers remain to be discovered in plants.

---

Plant life is exposed to potential threats from reactive oxygen species (ROS). During photosynthesis in plants, singlet oxygen (^1^O_2_) and superoxide radical (O_2_^−^) are constantly produced in chloroplasts (Asada 2006). Furthermore, the H_2_O_2_ generation flux in peroxisomes in illuminated leaves of C3 plants reaches 25% of the flux of CO_2_ fixation on electron base. This is approximately 50 times greater than the ROS generation flux associated with mitochondrial respiration (Foyer and Noctor, 2003). When plants suffer environmental stress, i.e. any changes in the surrounding environmental conditions that are not favorable for a plant, ROS production in the cells are increased due to the disturbance of metabolism (for example, the rate of CO_2_ fixation in leaves decreases at low temperatures or under drought). ROS oxidize biomolecules such as proteins, nucleic acids and lipids, thereby impairing their functions and damaging cells. As a defense system against ROS, plant cells are richly equipped with multiple types of ROS-scavenging enzymes and antioxidants, thereby keeping ROS concentrations low enough.

ROS exert their biological functions also via the secondary products ‘reactive carbonyl species (RCS)’. RCS is a collective name of the α,β-unsaturated aldehydes and ketones such as acrolein (Acr) and 4-hydroxy-(*E*)-2-nonenal (HNE) that are formed via the degradation of lipid peroxides (Esterbauer et al. 1991, Aldini et al. 2005, Mano 2012). An RCS molecule readily forms an adduct with a protein at the thiol-, amino-, or imidazole group via Michael addition of its β-carbon (Esterbauer et al. 1991), and cause alterations of the protein function (Wible and Sutter 2017). In plants, more than a dozen types of RCS occur (Schauenstein et al. 1977, Mano 2012) and their levels are increased under oxidative stress conditions (Yin et al. 2010, Yamauchi et al. 2012, Biswas and Mano 2015, Roach et al. 2018, Sultana et al. 2022, 2024). Rises of RCS levels have significant physiological impacts. For example, the genetic enhancement of the RCS-scavenging enzyme 2-alkenal reductase (AER) (Mano et al. 2002, 2005) in transgenic plants suppressed the increases in RCS levels due to salt stress (Sultana et al. 2024), aluminium stress (Yin et al. 2010) and light stress (Mano et al. 2010) and thereby conferred stress tolerance. Vice versa, the genetic deficiency of aldehyde alkenal/one reductase (Yamauchi et al. 2012) and aldehyde oxidase (Srivastava et al. 2017, Nurbekova et al. 2021) made the plants more susceptible to oxidative stress or senescence.

The plant defense system against RCS toxicity consists of several types of enzymes and small molecules, as the system against ROS. As for enzymatic defense, several classes of enzymes with distinct RCS-detoxifying reactions have been characterized such as AER (Mano et al. 2002, 2005), glutathione transferase tau isozymes (Mano et al. 2017, 2019a,b) and aldehyde-scavenging enzymes such as aldo-keto reductase (Yamauchi et al. 2011), aldehyde dehydrogenase (Sunker et al. 2003) and aldehyde oxidase (Srivastava et al. 2017, Nurbekova et al. 2021).

RCS-scavenging small molecules also occur in plants. Thiol compounds such as cysteine (Cys) and the reduced form of glutathione are excellent RCS scavengers (Esterbauer et al. 1975). Various polyphenols also have the RCS scavenging ability. Zhu et al. (2009) compared the RCS scavenging ability of a dozen tea polyphenols *in vitro* and found that nine of them, including epigallocatechin gallate (EGCG), showed a high Acr-scavenging activity. The reaction of EGCG and Acr starts with a nucleophilic attack of the carbon at position 8 on the EGCG A-ring on the *β*-carbon of Acr, thus forming a Michael adduct. Subsequent aldol condensation produces a ring structure (Huang, et al., 2020). One EGCG molecule can bind up to three Acr molecules (Sugimoto, et al., 2021). Acr-scavenging reactions have been also characterized for other polyphenols and their glycosides such as resveratrol, hesperetin (Wang, et al., 2015), ferulic acid (Tao, et al., 2019), pelargonidin (Colzani, et al., 2018), myricetin (Zhang, et al., 2020) and cyanidin-3-*O*-glucoside (Song, et al., 2021).

Amino compounds also have RCS-scavenging potential. The primary amino group in γ-aminobutyric acid (GABA), L-Ala and L-Ser serves as a Michael acceptor to bind up to two molecules of Acr (Jiang et al. 2020a, Zou et al. 2021). Recently it was reported that *S*-allyl cysteine sulfoxide (alliin), an amino acid typical of *Allium* species, can scavenge Acr at a high speed although the reaction mechanism has yet to be clarified (Uemura et al. 2023).

Theophylline, a purine derivative found in tea, forms a Michael adduct between its secondary amine and the *β*-carbon of Acr at a high rate (Jiang, et al., 2020b). Interestingly, animals contain RCS-scavenging dipeptides carnosine, homocarnosine, and anserine (Aldini et al. 2002) in brain and muscle. The amino and imidazole groups of these histidine-containing dipeptides react as Michael acceptors (Aldini et al., 2005), allowing them to be excellent RCS scavengers (Aldini et al., 2002; Liu et al., 2003; Carini et al., 2003). Exogenous application of these dipeptides to plant cells or seedlings suppressed the RCS levels and mitigated oxidative injury, without affecting ROS levels (Biswas and Mano, 2015; Sultana et al., 2022), showing that RCS-scavenging compounds can protect plants as the “secondary antioxidants”. In this study, we aimed at discovering RCS-scavenging compounds in plants. Employing Acr as a representative RCS, we evaluated RCS-scavenging ability of dozens of angiosperms.

Garlic cloves showed the highest content of Acr-scavenging ability, which was attributed to alliin, a garlic-typical amino acid. We found that isoalliin and methiin, the cysteine derivatives relative to alliin, also showed excellent Acr-scavenging ability and revealed the reaction mechanism. Potential of plant specialized compounds as the secondary antioxidants is discussed.

## Materials and Methods

### Chemicals

Alliin and methiin were purchased from LKT Laboratories (Saint Paul, MN, USA). Carnosine, fluorescein, and HNE dimethyl acetal (HNE-DMA) were purchased from Sigma-Aldrich Japan (Tokyo, Japan), and isoalliin from Nagara Science Co. (Gifu, Japan). Anserine nitrate was purchased from Fujifilm Wako Pure Chemical (Osaka, Japan). 2,4-Dinitrophenylhydrazine (DNPH), purchased from Fujifilm Wako, was purified by recrystallization in acetonitrile and kept in acetonitrile until use (Mano et al. 2022). All other chemicals were of analytical grade. Acr was prepared on the day of use by hydrolysis of the acetal, as follows. Acrolein dimethyl acetal (Tokyo Chemical Industry, Tokyo, Japan) was dissolved in 100 mM HCl aqueous solution at 40°C for 40 min. The solution was then neutralized with NaHCO_3_ and further incubated at 25°C for 2 h. The concentration of Acr was determined from the absorbance at 215 nm with an extinction coefficient of 15.0 mM^−1^ cm^−1^ (in water).

### Plant Materials

Bamboo (*Phyllostachys heterocycla*) shoots and yuzu (*Citrus junos*) fruits were harvested from Yamaguchi University’s experimental field. Following materials of fresh plants were purchased from grocery stores in Yamaguchi (Japan) between April and December 2018: Apple (*Malus domestica*) fruits, asparagus (*Asparagus officinalis*) shoots, avocado (*Persea americana*) fruits, bell pepper (*Capsicum annuum* ‘Grossum’) fruits, bitter melon (*Momordica charantia* var. *pavel*) fruits, broccoli (*Brassica oleracea* var. italica) buds, carrot (*Daucus carota* subsp. *sativus*) roots, cherry (*Cerasus avium*) fruits, chestnuts (*Castanea crenata*), Chinese yam (*Dioscorea polystachya*) roots, common ginger (*Zingiber officinale*) rhizomes, edible burdock (*Arctium lappa*) roots, eggplant (*Solanum melongena*) fruits, fig (*Ficus carica*) fruits, garlic (*Allium sativum*) cloves, grape (*Vitis labruscana* ‘Pione’) fruits, green onion (*Allium fistulosum*) leaves, guava (*Psidium guajava*) fruits, Haskap berry (*Lonicera caerulea* var. *emphyllocalyx*) fruits, Indian gooseberry (*Phyllanthus emblica*) fruits, Japanese apricot (*Prunus mume*) fruits, Japanese butterbur (*Petasites japonicus*) stalks, Japanese ginger (*Zingiber mioga*) shoots, Jun-sai (*Bransenia shreberi*) leaves, Kaki persimmon (*Diospyros kaki*) fruits, kiwifruit (*Actinidia deliciosa*) fruits, Korean lettuce (*Lactuca sativa*) leaves, loquat (*Rhaphiolepis bibas*) fruits, lotus (*Nelumbo nucifera*) rhizomes, Mandarin orange (*Citrus unshiu*) fruits, mulkhiya (*Corchorus olitorius*) leaves, okra (*Abelmoschus esculentus*) fruits, onion (*Allium cepa*) bulbs, parsley (*Petroselinum crispum*) leaves, pea (*Pisum sativum*) seeds, perilla (*Perilla frutescens* var. *crispa*) leaves, pitaya (*Hylocereus undatus*) fruits, plum (*Prunus salicina*) fruits, spinach (*Spinacia oleacea*) leaves, star fruit (*Averrhoa carambola*) fruits, strawberry (*Fragaria* × *ananassa*) achenes, taro (*Colocasia esculentai*) tubers, watermelon (*Citrullus lanatus*) fruits, and white radish (*Raphanus sativus* var. *hortensis*) roots.

### Plant Extract Preparation

An edible portion (10 g) taken from a fresh sample was homogenized in 80% (v/v) ethanol (20 mL) using a kitchen blender and filtered through a layer of Miracloth (Sigma-Aldrich Japan). The filtrate was acidified with HCl (10 mM final) to facilitate protein precipitation. It was incubated at 25°C for 10 minutes, centrifuged at 9,000 × *g* for 30 min, and the resulting supernatant was neutralized with sodium bicarbonate. The volume of deproteinized extract was measured.

### Determination of the Acr-scavenging ability

A reaction mixture containing 200 μM Acr, 50 mM potassium phosphate pH 6.8, and a plant extract was incubated at 30°C for 10 min. One volume of the mixture was then added to 6.35 volumes of acetonitrile containing 4 mM DNPH and 0.4 M formic acid and incubated at 30°C for 60 min. The resulting hydrazone solution was filtered through a 0.45 μm filter (Millex LH; Merck, Darmstadt, Germany) and analyzed by HPLC, as follows. The hydrazone solution (10 μL) was injected into a Zorbax ODS column (4.6 × 250 mm, 5 μm; Agilent Technologies Japan, Hachioji, Japan) kept at 40°C and separated in a 1.0 mL min^−1^ flow of 75% (v/v) acetonitrile in water (isocratic elution) with a high-pressure pump (2690 Separation Module, Waters Corp., Milford, MA). The hydrazone was detected with a UV-visible detector (2487 Absorbance Detector, Waters) at 370 nm. This concentration was chosen based on reported values in Arabidopsis seedlings under salt stress (Sultana et al. 2022), corresponding to ∼200 µM tissue concentrations, thereby approximating physiologically relevant stress levels.

The decrease rate (*v*) of Acr concentration in the reaction mixture (in mol Acr consumed s^−1^ L^−1^) was used to calculate the Acr-scavenging ability of the plant extract per unit volume (in mol Acr s^−1^ (L extract)^−1^) by multiplying *v* by the volume ratio *P* (reaction mixture/plant extract) as follows:

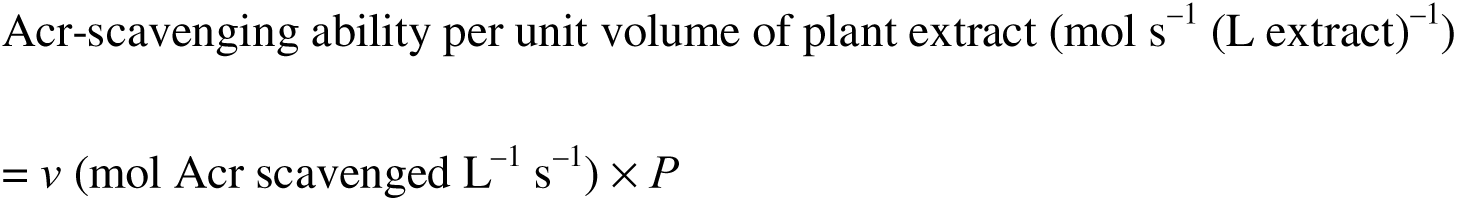

The Acr-scavenging ability content in the original sample, as mol Acr consumed s^−1^ (g sample)^−1^, was then calculated from the extraction efficiency *E* (L extract (g sample)^−1^) as follows:

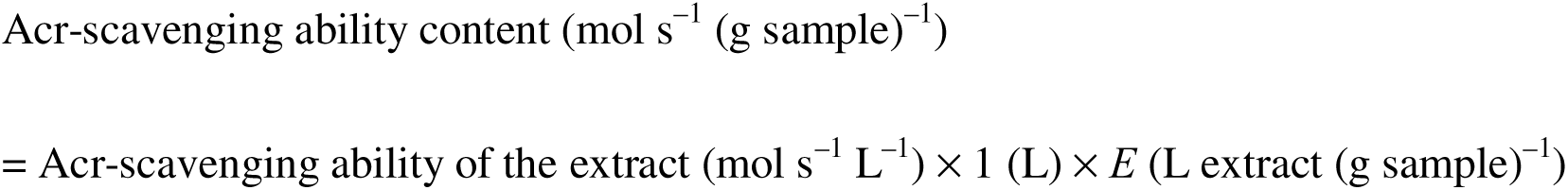

### Purification of the Acr-scavenging component from garlic cloves

Garlic cloves (40 g) were homogenized in 80% (v/v) ethanol (80 mL) using a kitchen blender, filtered through two layers of Miracloth, and centrifuged at 9,000 × *g* for 30 min to remove insoluble debris. The supernatant was freeze-dried to obtain 7.6 g of material, which was dissolved in distilled water (15 mL) and loaded onto a preparative hydrophilic column (Universal Column Premium Amino 30 μm, 3.0 × 16.5 cm; Yamazen Corp., Osaka, Japan) with a medium pressure chromatography system (EPCLC AI-580; Yamazen Corp.) equipped with a detector (fixed wavelength at 254 nm). Chromatography was performed at a flow rate of 20 mL/min under the following solvent conditions: solvent A: acetonitrile:10 mM ammonium acetate in water (88:12, v/v); solvent B: acetonitrile:100 mM ammonium acetate in water (1:1, v/v). The gradient program consisted of 0–16 min isocratic elution of 0% B, 16–56 min of a linear gradient from 0% to 100% B, and 56–80 min isocratic elution of 100% B. The fraction size was 20 mL. Each fraction was freeze-dried and then dissolved in 2 mL of distilled water, and its Acr-scavenging ability was determined as described above. The most active fraction was then separated on a Unison UK-Amino Column (3 μm, 4.6×100 mm; Imtakt, Kyoto, Japan) with an HPLC apparatus equipped with an LH-40 liquid handler and an SPD-10AVP UV-VIS detector (Shimadzu Corp., Kyoto, Japan). Chromatography was performed at a flow rate of 0.6 mL min^−1^ with the following multistep gradient elution: 0–10 min isocratic elution of 0% B; 10–35 min of a linear gradient from 0% to 100% B; 35–50 min isocratic elution of 100% B; 50–55 min of a linear gradient from 100% to 0% B; and, finally, 55–90 min isocratic elution of 0% B. The fraction size was 1.8 mL. The elution profile was monitored at 220 nm for nonspecific detection of organic substances. After 40 separation runs (50 μL injection per run), identical fractions were accumulated, freeze-dried, and dissolved in a smaller volume to determine the Acr-scavenging ability.

### LC-MS/MS analysis

Chromatographic separation was performed on a hydrophilic Intrada Amino Acid column (3 µm, 3 × 100 mm; Imtakt) with a Vanquish UHPLC System (Thermo Fisher Scientific, Waltham, MA). A 5-μL aliquot of the sample was injected into the column and eluted at a flow rate of 0.4 mL min^−1^ under the following solvent conditions: solvent A: 0.3% (v/v) in acetonitrile; solvent B: 0.1 M ammonium formate in water:acetonitrile (80:20, v/v). The gradient program was as follows: 0–10 min isocratic elution of 15% B; 10–25 min of a linear gradient from 15% to 60% B; 25–40 min isocratic elution of 60% B; 40–45 min of a linear gradient from 60% to 15% B; and finally 45–50 min isocratic elution of 15% B. The column temperature was set to 40°C. Mass spectra were acquired using an Orbitrap Exploris 120 mass spectrometer (Thermo Fisher Scientific) equipped with an H-ESI source, with acquisition in positive ion mode. The source parameters were as follows: capillary voltage of 3.5 kV in positive ion mode, ion transfer tube temperature of 325°C, vaporizer temperature of 350°C, sheath gas flow of 50 units, auxiliary gas flow of 10 units, and sweep gas of 1 unit. A full scan was performed in the mass-to-charge ratio (*m*/*z*) range of 100–1000, with a resolution of 120,000 at *m*/*z* 200. Tandem MS information was acquired in tMS^2^ mode with a resolution of 12,000 and HCD collision energies of 15, 30, and 45 eV.

## Results and Discussion

### Comparison of Acr-scavenging ability of plants

For choosing a starting material suitable for purifying RCS-scavenging compounds, we first collected various plant samples across diverse angiosperm lineages. To obtain samples in such a variety in short time, we obtained them (46 species in total), mainly fruits and vegetables, from local retail stores (see Materials and Methods). Because polyphenols have been extensively investigated as acrolein scavengers (Colzani, et al., 2018; Huang, et al., 2020; Sugimoto, et al., 2021; Tao, et al., 2019; Wang, et al., 2015; Zhang, et al., 2020; Zhu, et al., 2009), we aimed at polar compounds in this study. We chose 80% ethanol as an extraction medium, which is preferable to extracting more polar compounds than polyphenols (Phahom and Mano, 2022). This extract was deproteinized by acid treatment, then neutralized, and its Acr-scavenging rate was determined. To survey RCS scavengers that can react rapidly with Acr, we determined Acr consumption in the initial 10 minutes in our assay. The initial Acr concentration was set at 200 μM, mimicking physiological concentration, which was estimated on the following observation. A typical Acr content in salt-stressed *A. thaliana* seedlings is 200 nmol (g fresh weight)^−1^ (Sultana, et al., 2022). From this value, Acr concentration of 200 μM in the tissue was calculated on a simple assumption that the Acr molecules occur in the tissue in a homogenous solution of 1 mL volume per gram tissue weight. The reaction temperature was set at 30°C to simulate field conditions.

Fig. 1 summarizes the Acr-scavenging ability on the fresh weight base of the plant materials. Most plant materials showed significant levels of Acr-scavenging ability, thus indicating that many plants contain Acr-scavenging compounds. The highest Acr-scavenging ability content (in nmol Acr scavenged s^−1^ (g fresh weight)^−1^) were 1.39 for garlic cloves, 0.99 for spinach leaves, 0.96 for avocado fruits, 0.92 for lotus rhizomes, 0.91 for broccoli buds, 0.82 for chestnuts, 0.70 for bamboo shoots, 0.67 for onion bulbs, and 0.65 for both Chinese yam roots and white radish roots. These plants, when placed in the angiosperm phylogeny tree (Cole, et al., 2019), were distributed to different orders of taxa (Fig. 1). We chose garlic cloves as the starting material for purification to identify an Acr-scavenging compound.

**Fig 1.**
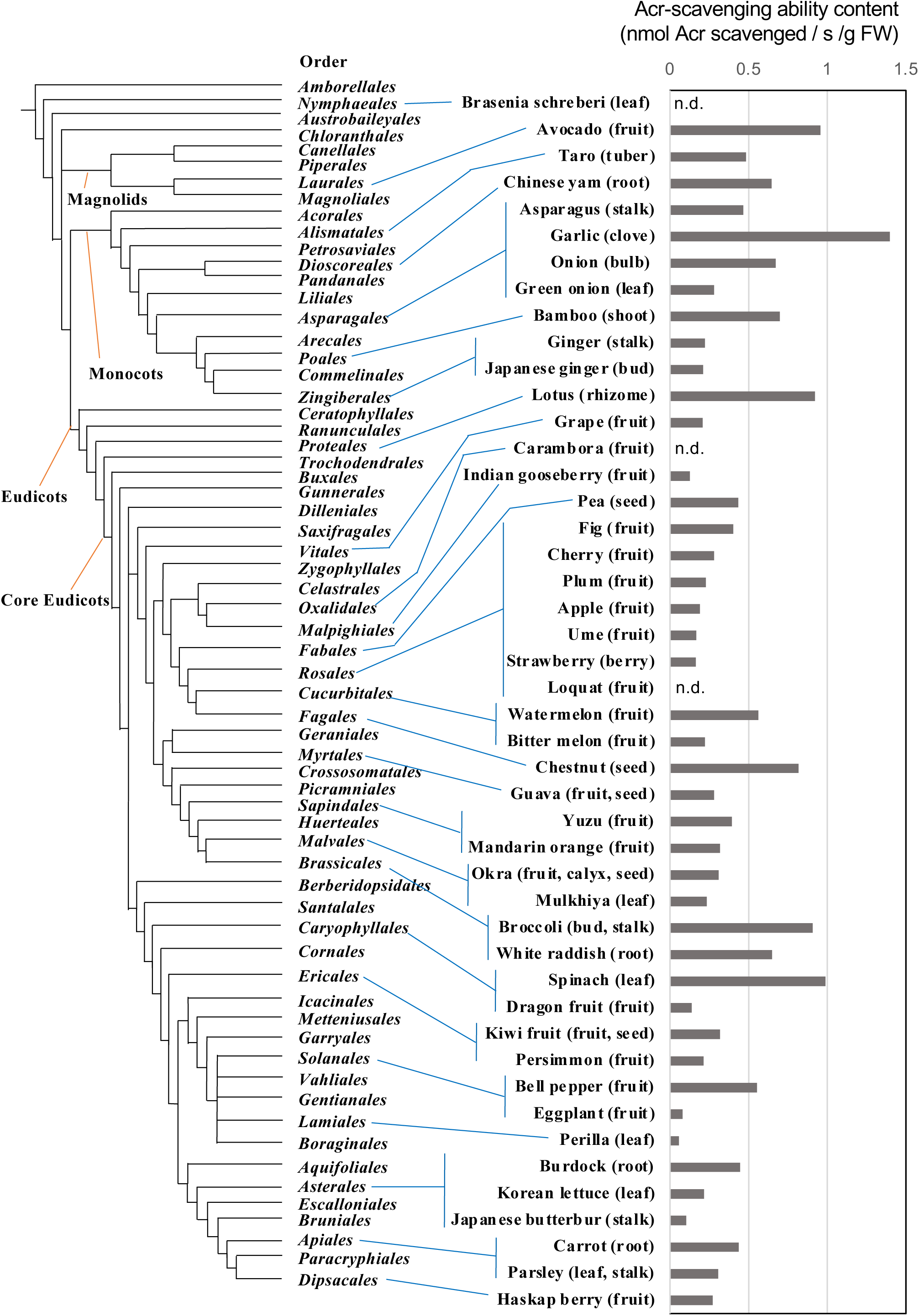
Acr-scavenging ability contents of plant materials. Plant extracts (80% ethanol) were deproteinized, neutralized, and tested for the Acr-scavenging ability as in Materials and Methods. Data of two runs (white dots) and their average (gray bar) is shown. The phylogenic tree was constructed using the Angiosperm Phylogeny Poster (Cole et al. 2019) as a guide.

### Purification of Acr-scavenging components from garlic extract

Garlic clove extract was first separated by preparative amino column chromatography (Fig. 2), which showed a single prominent peak of Acr-scavenging ability in fractions #13, #14, and #15. We scaled up the sample amount and the column size, as described in the Materials and Methods section, collected the active fractions, concentrated and purified them using an analytical HPLC column. The resulting fraction #8 had a high Acr-scavenging ability (Supplemental Fig. S1). We then analyzed this fraction by LC-MS/MS. Total ion chromatography (*m*/*z* 100–1000) revealed three major components: a broad peak eluted at 15.42 min with *m*/*z* 178.0528, a smaller peak at 16.13 min with *m*/*z* 360.1495 and a narrow peak at 18.87 min with *m*/*z* 258.1096. Mass spectra of these peaks are shown in Fig. 3A.

**Fig. 2.**
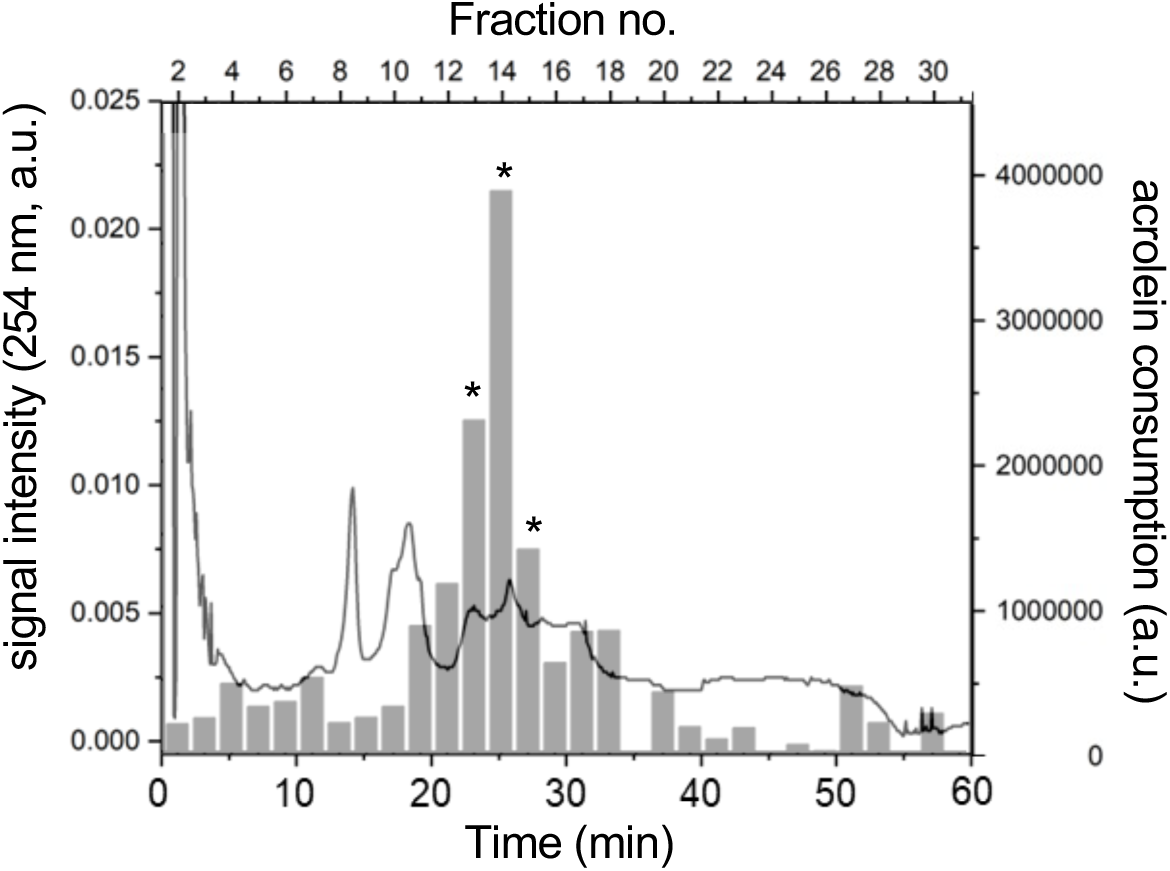
A typical result of the first step of purification of Acr-scavenging substances from garlic extract. Garlic extract was freeze-dried, dissolved in distilled water, applied onto a Premium Amino Column (Yamazen) and then separated as described in the Materials and Methods. Solid line, absorbance at 254 nm. Gray bars, Acr consumption by each fraction, determined as described in Materials and Methods. The active fractions (no. 13, 14 and 15, asterisks) were collected for further purification.

**Fig. 3.**
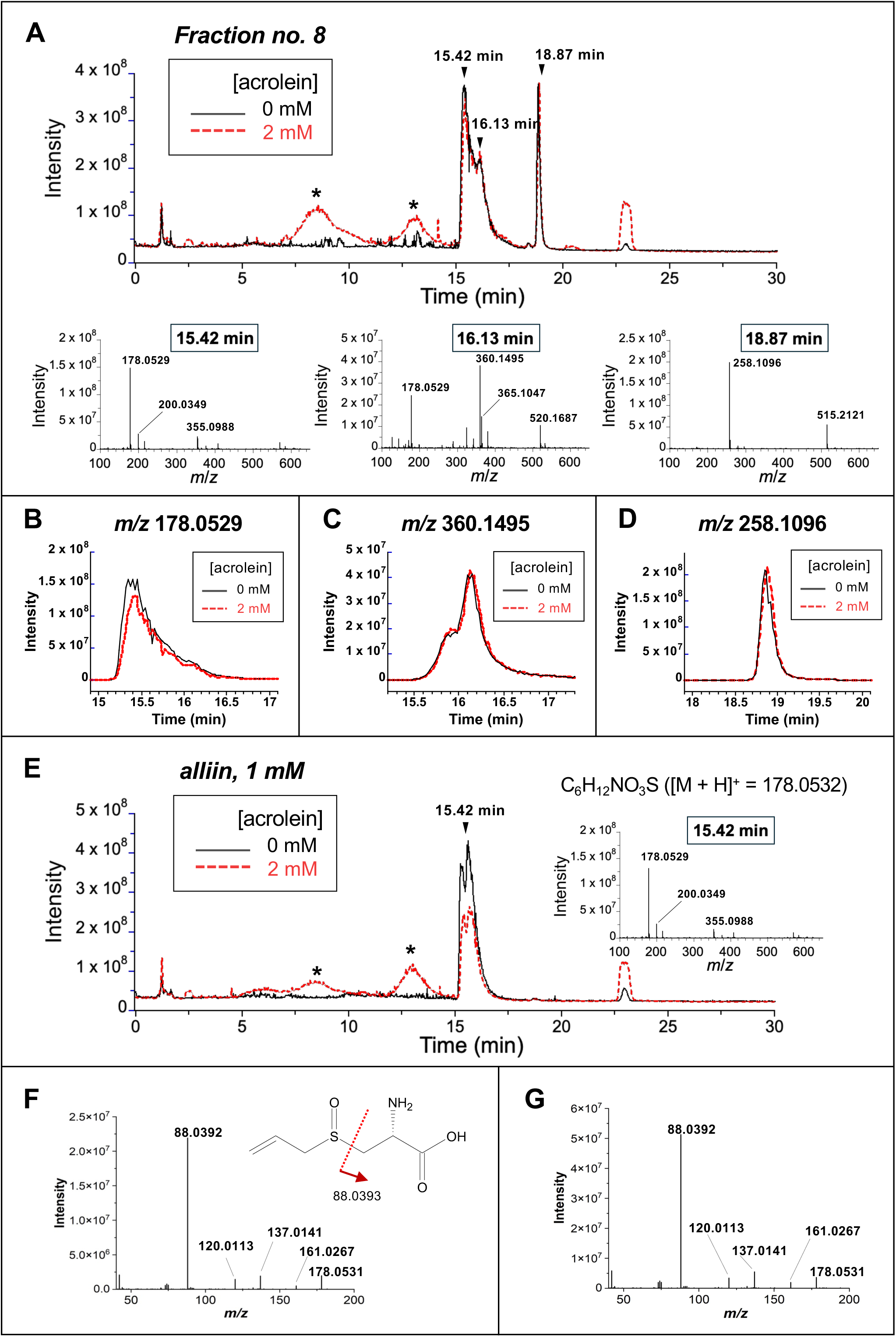
LC-MS/MS evidence that alliin is the Acr-scavenging substance in fraction #8. A: Total ion chromatograms (*m*/*z* range 100–1000) of fraction #8 obtained from the second step of purification (Supplementary Fig. S1) with 2 mM Acr (red broken trace) and without it (black solid). Asterisks show possible reaction products of the scavenging compound with Acr. The MS spectra of the signals at 15.42 min, 16.13 min and 18.87 min are shown below the chromatogram. No significant signals were detected in the *m*/*z* range over 650. B, C and D: Ion chromatograms of the *m*/*z* 178.0529 signal (*m*/*z* range 178.0520-178.0538), m/z 360.1495 (*m*/*z* range 360.1478-360.1514) and *m*/*z* 258.1096 (*m*/*z* range 258.1082-258.1108), respectively, extracted from the Fig. 3A data, with 2 mM Acr and without it. E: Total ion chromatograms (*m*/*z* range 100–1000) of 1 mM alliin with 2 mM Acr (red broken trace) and without it (black solid). F: The MS/MS spectrum of the *m*/*z* 178.0532 signal of alliin. The fragmentation position corresponding to the m/z 88.0932 signal is shown in the formula. G: The MS/MS spectrum of the *m*/*z* 178.0532 signal at 15.42 min in fraction #8.

To clarify which component is the Acr scavenger, we mixed fraction #8 with Acr and examined the changes in the total ion chromatogram (Fig. 3A, broken trace in red). The addition of Acr at 2 mM resulted in a significant decrease in the compound with *m*/*z* 178.0529 peaking at 15.42 min (Fig. 3B), while those with *m*/*z* 360.1485 at 16.13 min (Fig. 3C) and *m*/*z* 258.1096 at 18.87 min (Fig. 3D) were unchanged, indicating that the *m*/*z* 178.0528 signal represented the Acr scavenger in fraction #8. Addition of Acr to fraction #8 generated two new broad peaks at 8.57 (*m*/*z* 290.1056) and 13.16 min (*m*/*z* 234.0795) (Fig. 3A, asterisks). The *m*/*z* values of these peaks matched hypothetical Acr adducts to the *m*/*z* 178.0528 compound; the 13.16 min peak corresponding to a mono-Acr adduct (mass increment 56.0265, matching the molecular weight of Acr (C_3_H_4_O)), and the 8.57 min peak a di-Acr adduct (mass increment 112.0526).

Assuming that the *m*/*z* 178.0528 ion represents a [M + H] ^+^ ion, the elemental composition of the molecule was deduced as C_6_H_11_NO_3_S. Retrieval of the corresponding compounds in the databases AraCyc (Mueller, et al., 2003), PlantCyc (Plant Metabolic Network (PMN): https://pmn.plantcyc.org/organism-summary?object=PLANT, on www.plantcyc.org, Jan 15, 2022), and KEGG (Kanehisa, et al., 2000) using the software package Compound Discoverer 3.0 (Thermo Fisher Scientific), resulted in *S*-allyl-cysteine sulfoxide (alliin; C_6_H_11_NO_3_S, 177.22 g mol^−1^) as the most probable candidate.

Alliin is an amino acid sulfoxide characteristic of garlic, accounting for about 30% (w/w) of total free amino acids in garlic cloves (Ueda, et al., 1991), and is a precursor of allicin, the major compound of garlic odor (Stoll and Seebeck, 1948). In the LC-MS/MS analysis, authentic L-alliin eluted as a broad peak ranging from 15.2–16.7 min with an *m*/*z* of 178.0529 (Fig. 3E). The MS^2^ spectrum of this parent ion (Fig. 3F) matched the spectrum of the *m*/*z* 178.0529 ion of the Acr scavenger in fraction #8 (Fig. 3G). The addition of Acr to alliin decreased the alliin peak and generated two broad peaks at 8.20 (*m*/*z* 290.1056) and 13.02 min (*m*/*z* 234.0795) (Fig. 3E), as observed for fraction #8 (Fig. 3A). Thus, the Acr-scavenging component in the purified fraction #8 was identified as alliin.

Alliin appears to be the main component responsible for the Acr-scavenging activity in garlic extract. This conclusion is based on the following estimation. As shown in Fig. 2A, alliin accounted for most of the Acr-scavenging activity in fraction #8. The scavenging activity of fraction #8 represented 24% of the total activity separated in the second chromatography step (Supplementary Fig. S1), in which the major activity (approximately 50%) was recovered from the original garlic extract during the first chromatography step (Fig. 2). Assuming that no activity was lost during the purification processes, alliin accounts for at least 12% of the total Acr-scavenging activity in the garlic extract.

### Reaction mechanism of alliin with Acr

Uemura et al. (2023) have reported that alliin has a high capacity to scavenge Acr. Its reaction mechanism with Acr, which remains to be elucidated, could be similar to that of an amino compound such as carnosine and GABA (Carini et al. 2003; Jiang et al. 2020a), as illustrated in Fig. 4. Specifically, the Michael addition of the *β*-carbon to the amino nitrogen, and the second addition of Acr to the same nitrogen, followed by the intramolecular aldol condensation to form a nitrogen-containing heterocycle. When alliin was incubated with Acr, new peaks appeared at 8.20 min and 13.02 min, with MS signals at *m*/*z* 234.0795 and *m*/*z* 290.1056, respectively. The intensities of both signals increased with increasing Acr concentration (Fig. 5A, B). The former MS signal corresponds to the mono-Acr adduct and the latter di-Acr adduct. In addition, another signal at *m*/*z* 272.0950 appeared at higher Acr concentration (Fig. 5C). On the assumption that it represented a [M + H]^+^ ion, the atomic composition of the compound was deduced as C_12_H_17_NO_4_S. We presumed that this signal corresponded to a dehydrated product of the di-Acr adduct.

**Fig. 4.**
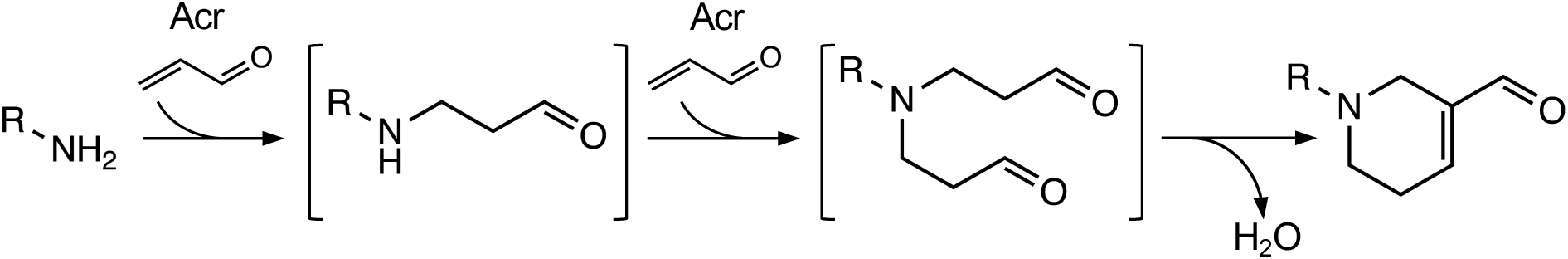
Sequential reactions of an amino compound with Acr molecules and subsequent dehydration

**Fig. 5.**
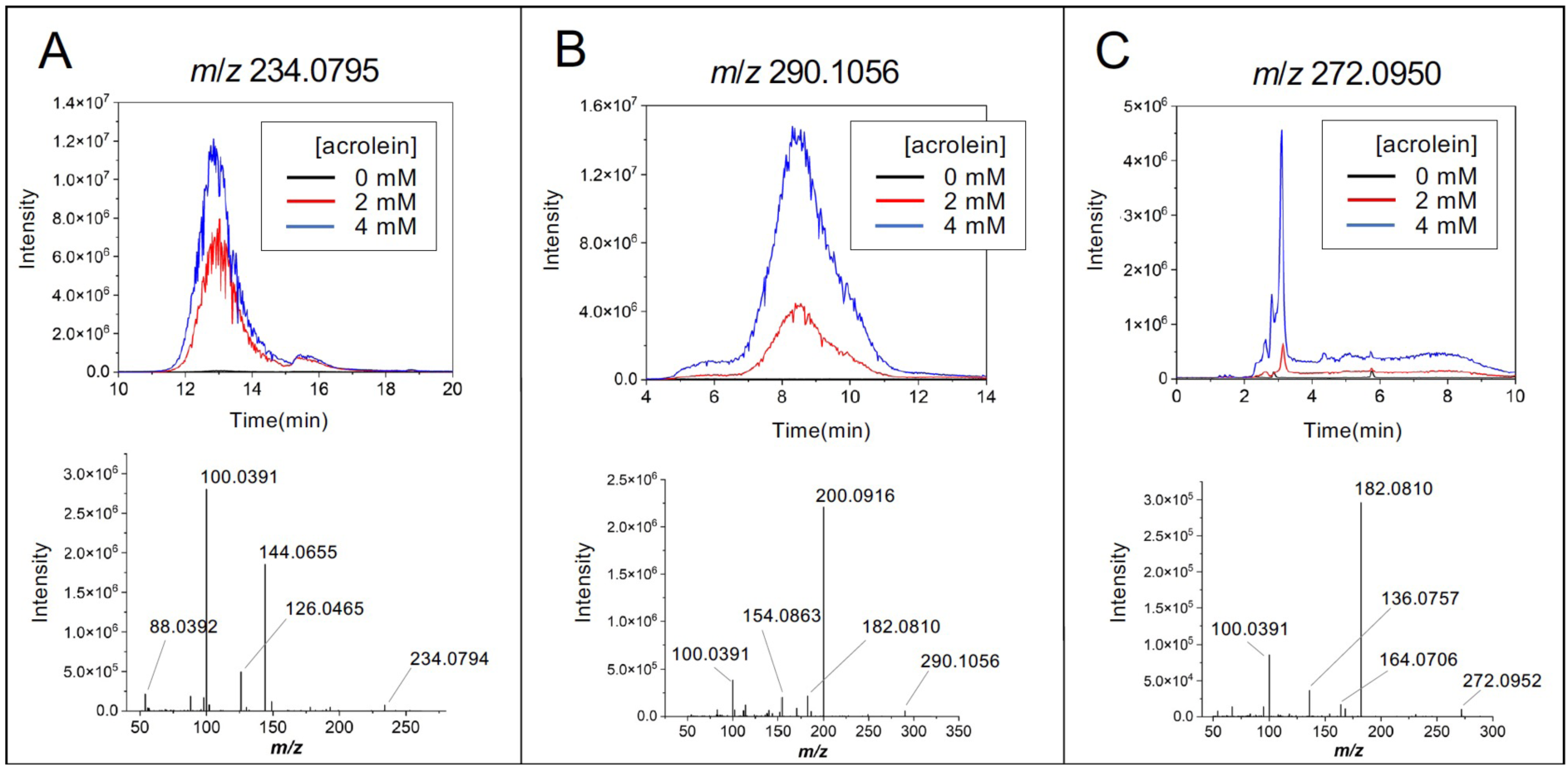
LC-MS/MS results for alliin-Acr adducts. An authentic alliin preparation (5 mM) was mixed with various concentrations of Acr, incubated for 1 h, and subjected to LC-MS/MS analysis. A: Extracted ion chromatograms of the mono-Acr adduct of alliin (*m*/*z* 234.0795). B: Extracted ion chromatograms of the di-Acr adduct (*m*/*z* 290.1056). C: Extracted ion chromatograms of the dehydrated products from the di-Acr adduct (*m*/*z* 272.0950). The corresponding parent ion MS/MS spectrum is shown at the bottom of panels A–C.

To elucidate the mechanism of the reaction between alliin and Acr, we purified the putative dehydrated product of the di-Acr adduct and determined its structure by NMR. The deduced structure corresponded to the *N*^b^-(3-formyl-3,4-dehydropiperidino) derivative of alliin (Fig. 6), which matches the di-Acr adduct expected from the reaction scheme illustrated in Fig. 4. Specifically, one Acr molecule initially adds to the amino group via a Michael-type reaction, and the resulting secondary amino group subsequently reacts with another Acr molecule to form the di-Acr adduct, which then undergoes intramolecular aldol condensation to yield a nitrogen-containing heterocycle. The MS² spectra support this reaction sequence: the mono-Acr adduct produced a fragment ion at *m*/*z* 144.0655, whose deduced composition (C[H[[NO[) corresponded to cleavage between the sulfur atom and the β-carbon of the amino acid (Fig. 6), identical to the cleavage observed for alliin (Fig. 3F). Likewise, the di-Acr adduct and its dehydrated product yielded fragment ions at *m*/*z* 200.0917 (Fig. 5B) and *m*/*z* 182.0811 (Fig. 5C), respectively, both consistent with the same cleavage position in the proposed structures.

**Fig. 6.**
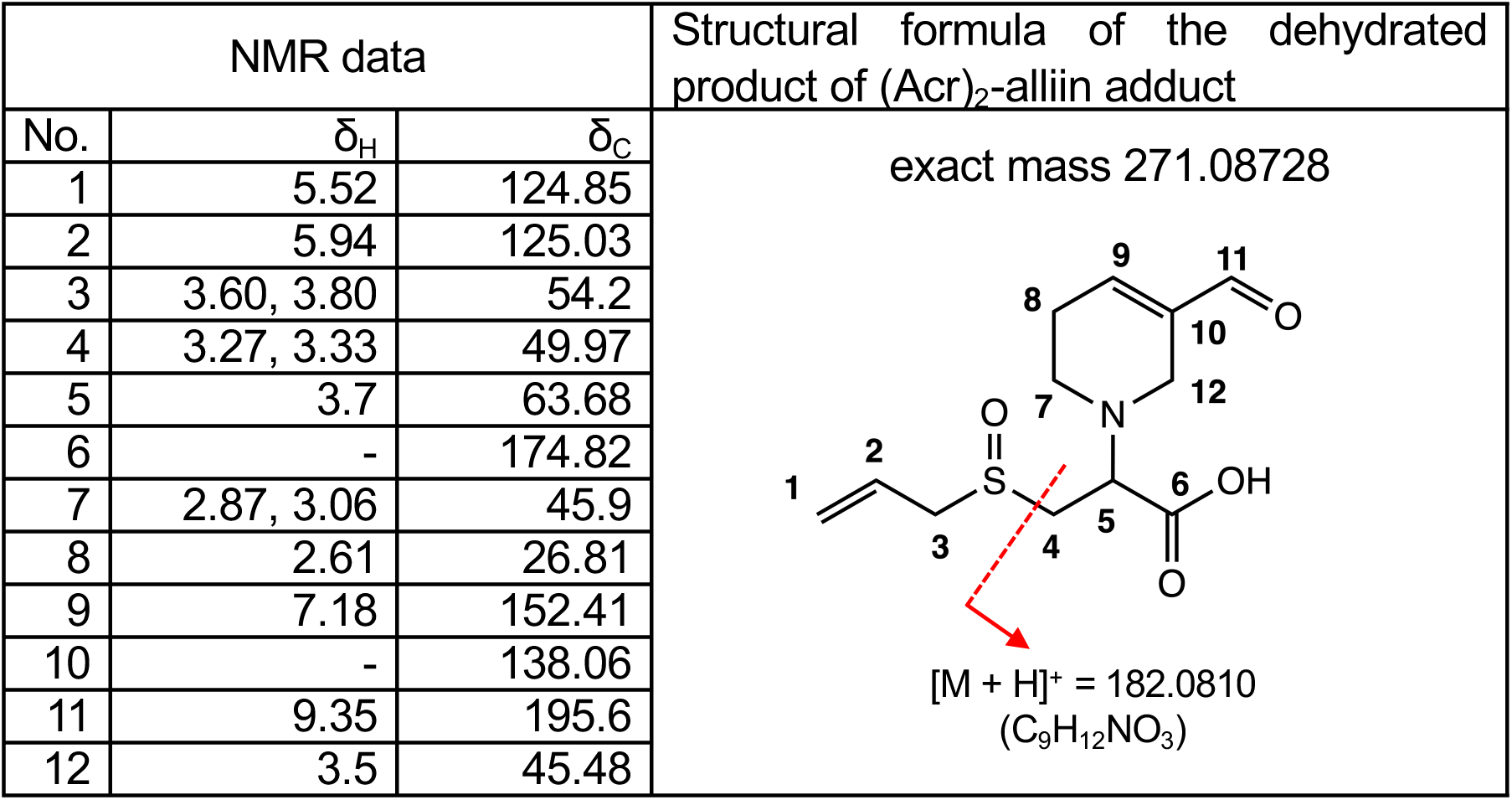
Structural analysis of the purified adduct of *m*/*z* 272.0950 with ^1^H NMR (400 MHz) and ^13^C NMR (500 MHz). Chemical shift data and the deduced structure of the dehydrated product of (Acr)_2_-alliin adduct are shown. Red broken line indicate the position of fragmentation that can generate *m*/*z* 182.0810 signal.

### Comparison of the Acr-scavenging ability of S-alk(en)yl cysteine sulfoxides, amino compounds, and polyphenols

Plants in the *Allium* family are rich in alliin-analogous *S*-alk(en)yl cysteine sulfoxides, such as *S*-propenylcysteine sulfoxide (isoalliin) in onion (*A. cepa*) and *S*-methylcysteine sulfoxide (methiin) in Oriental garlic (*A. tuberosum*) (Yamazaki, et al., 2011). Isoalliin and methiin also formed mono-Acr and (Acr)_2_ adducts and the dehydration product of the (Acr)_2_ adduct, as did alliin (Supplementary Fig. S2 and S3). The Acr-scavenging rates of these *S*-alk(en)yl cysteine sulfoxides were 120–150 μM Acr in 30 min, when the reaction was started with 1 mM Acr and 1 mM scavenger (Table 1A). These rates were comparable to, or even higher than, those for carnosine and anserine, Acr-scavenging dipeptides in animals (Carini, et al., 2003; Spaas, et al., 2021). As compared with broader variety of amino compounds, these *S*-alk(en)yl cysteine sulfoxides excelled GABA and most of protein-constituting amino acids such as Met, His, Lys, Arg and Gly for the Acr-scavenging rate. Such high Acr-scavenging rate for *S*-alk(en)yl cysteine sulfoxides appears to be attributed to the presence of sulfoxide close to the amino carbon. Indeed, methionine sulfoxide, in which the sulfoxide is one carbon further away, showed a significantly faster Acr-scavenging rate than other amino acids lacking a sulfoxide although its rate was slower than *S*-alk(en)yl cysteine sulfoxides (Table 1A).

**Table 1.** (A) Comparison of Acr-scavenging rate of amino compounds. Acr (1 mM) was incubated with a scavenger (1 mM) in 50 mM potassium phosphate, pH 7.0, at 30°C for 30 min. The remaining Acr was determined by HPLC after derivatization with DNPH, as described in the Materials and Methods section. Acr consumption (concentration in the reaction mixture) in 30 min is shown. (B) Comparison of Acr-scavenging rate for alliin, polyphenols, and theophylline. Acr (1 mM) was incubated with a scavenger (1 mM) in 10% DMSO, 50 mM potassium phosphate, pH 7.0, at 30°C for 5 h. Acr consumption in 5 h is shown. The values shown are the average and standard error of three independent runs. Different letters in the Statistics column represent significantly different values (*P* < 0.05, Tukey’s test).

| **A** | Compound | Acr consumption ($\mu\text{M}$ in 30 min) | Statistics |
| --- | --- | --- | --- |
| | Isoalliin | $153.8 \pm 6.5$ | a |
| | Methiin | $144.5 \pm 14.0$ | ab |
| | Alliin | $126.8 \pm 5.4$ | bc |
| | Carnosine | $114.3 \pm 1.3$ | cd |
| | Anserine | $99.4 \pm 5.5$ | de |
| | Methionine sulfoxide | $86.0 \pm 8.3$ | e |
| | Histidine | $63.9 \pm 6.0$ | f |
| | Lysine | $59.5 \pm 8.9$ | f |
| | Phenylalanine | $57.5 \pm 0.9$ | f |
| | EGCG | $45.4 \pm 5.2$ | f |
| | Arginine | $44.7 \pm 6.3$ | f |
| | Histamine | $40.1 \pm 8.4$ | fg |
| | Methionine | $22.7 \pm 8.7$ | gh |
| | GABA | $10.6 \pm 5.9$ | h |
| | β-Alanine | $6.5 \pm 3.7$ | h |
| **B** | Compound | Acr consumption ($\mu\text{M}$ in 5 h) | Statistics |
| | Alliin | $419.1 \pm 3.0$ | a |
| | EGCG | $282.0 \pm 4.6$ | b |
| | Resveratrol | $31.8 \pm 1.5$ | c |
| | Hesperetin | $18.9 \pm 1.0$ | d |
| | Theophylline | $17.9 \pm 3.6$ | d |

We also compared the *S*-alk(en)yl cysteine sulfoxides with polyphenols that can scavenge Acr. Isoalliin, methiin and alliin outperformed EGCG in scavenging Acr (Table 1A). We note that assays of polyphenols (resveratrol, hesperetin) were conducted in 10% DMSO with longer incubation times due to low solubility and slower reaction kinetics (Wang, et al., 2015). Therefore, direct quantitative comparisons with amino acid sulfoxides should be made cautiously. To compare alliin with resveratrol and hesperetin, a separate experiment was performed in 10% DMSO solution and reaction time was set to 5 h (Table 2B). Under these conditions, alliin scavenged Acr faster than EGCG, resveratrol, and hesperetin. Theophylline, another type of Acr scavenger (Jiang, et al., 2020b) showed an Acr-scavenging rate comparable to that for hesperetin and much slower than that for EGCG and alliin (Table 1B).

### RCS scavenging compounds are present in many plant species

Because RCS are secondary toxic products derived from ROS, plants have evolved multiple enzymatic and non-enzymatic mechanisms to mitigate their harmful effects (Mano et al. 2019a,b). In this study, we established a method to evaluate the RCS-scavenging ability of plant extracts and demonstrated that *S*-alk(en)yl cysteine sulfoxides, amino acids characteristic of *Allium* family, are highly efficient Acr scavengers. Among them, isoalliin, the major sulfoxide in onion, exhibited the strongest Acr-scavenging activity, surpassing not only its structural analog alliin but also other well-known plant-derived and animal-derived scavengers. This finding identifies isoalliin as a representative plant RCS scavenger and underscores the contribution of *Allium* metabolites to the chemical defense system against carbonyl stress. Isoalliin concentrations in onion bulbs are reported at 5–20 µmol g[¹ FW (Yamazaki et al. 2011; Ueda et al. 1991). At these levels, isoalliin would be comparable to stress-induced Acr accumulation, supporting its role as an in planta scavenger.

It is noteworthy that the presence of a sulfoxide group in close proximity to the amino carbon markedly enhances the Michael acceptor reactivity of the amino group, thereby conferring the high RCS-scavenging capacity observed in S-alk(en)yl cysteine sulfoxides. Consequently, the RCS-scavenging property of a compound appears to be governed by the cooperative action of two or more functional groups, rather than by the reactivity of a single moiety alone. More broadly, our findings suggest that structurally diverse, and as yet unexplored, plant metabolites may play critical roles in mitigating oxidative damage through RCS detoxification.

## ABBREVIATIONS

Acr: acrolein
DNP-: dinitrophenylhydrazo-
DNPH: 2,4-dinitrophenylhydrazine
FW: fresh weight
GSH: the reduced form of glutathione
HNE: 4-hydroxy-(*E*)-2-nonenal
LOOH: lipid peroxide
RCS: reactive carbonyl species
ROS: reactive oxygen species

## AUTHOR CONTRIBUTION

Conceptualization: J. M. and D.S.; Investigation: A.H., N.T.; Methodology: J.M., C.N., A.H., N.T., Y.M., K.M.; Roles/Writing - original draft: A.H., N.T., Writing - review & editing; A.H., J.M., Funding acquisition: J.M., D.S.; Supervision: J.M.

## ASSOCIATED CONTENTS

Supporting information is available.

Chromatographic results for the purification of Acr-scavenging substances from garlic extract (Fig. S1), LC-MS/MS results for the reaction products of isoalliin and Acr (Fig. S2), and LC-MS/MS results for the reaction products of methiin and Acr (Fig. S3).

## ACKNOWLEDGMENTS

The authors would like to express their gratitude to Ryoma Oishi and Suzuka Monden for their technical assistance and to Prof. Toshihiro Murafuji, Yamaguchi University, for discussion.

## FUNDING

This work was supported by the Ministry of Agriculture, Forestry and Fisheries of Japan, by KAKENHI Grants (nos. 17K19909 and 20H03278) from the Japan Society for the Promotion of Science (to J. M.), and by Yamaguchi University Project for Formation of the Core Research Project. This work resulted from the use of research equipment shared in the MEXT Project for promoting public utilization of advanced research infrastructure (Program for supporting the construction of core facilities) Grant Number JPMXS0440400022.

## Conflict-of-Interest Declaration

The authors declare no conflicts of interest associated with this manuscript.

## Declaration of generative AI and AI-assisted technologies in the writing process

During manuscript preparation, the authors used AI-based language tools (DeepL, ChatGPT, Google Gemini) solely for improving English grammar and readability. All scientific content was conceived, verified, and revised by the authors, who take full responsibility for the manuscript.

## Deposition of a preprint

This manuscript has been deposited as a preprint at bioRxiv (DOI: 10.1101/2025.08.18.670978).

## Notes

### Competing Interest Statement

The authors have declared no competing interest.

### Summary of Updates

1) The title is changed to represent the content more properly. 2) One Figure is moved from Supplementary data to the main data. 3) The figure of the chromatographic evidence for the identity of the acrolein scavenger in garlic is revised so that it shoes more MS data. 4) Figures are renumbered.

