## Supplementary Figure S1 for "*S*-Alk(en)yl-Cysteine Sulfoxides in Allium Species Are Excellent Acrolein Scavengers: Implications for Secondary Antioxidants in Plants"

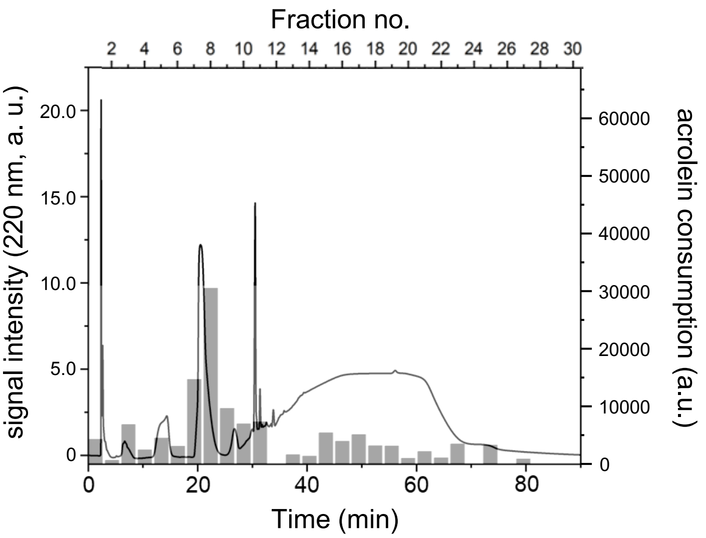


**Supplementary Figure S1.** A typical result of the second step of purification of Acr-scavenging substances from garlic extract. shown. Active fractions obtained from the first step (Fig. 2) were collected, freeze-dried, dissolved in distilled water and applied to a Unizon UK-Amino column (3 μm, 4.6×100 mm, Imtakt). Solid line, signal intensity at 220 nm. Gray bars, Acr consumption by each fraction.
