## Supplementary Figure S2 for "*S*-Alk(en)yl-Cysteine Sulfoxides in Allium Species Are Excellent Acrolein Scavengers: Implications for Secondary Antioxidants in Plants"

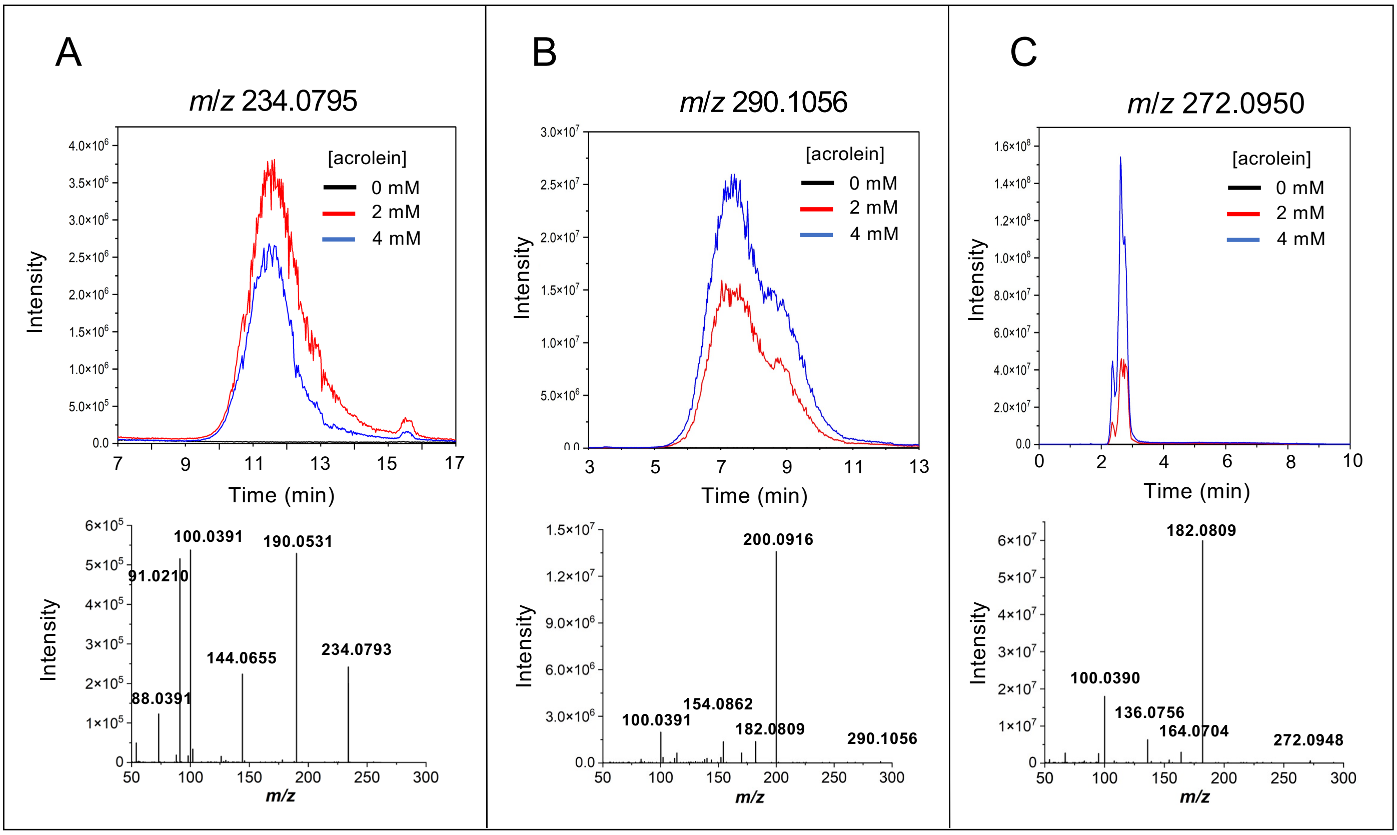


**Supplementary Figure S2.** LC-MS/MS analysis of the reaction products of isoalliin and Acr. An authentic isoalliin preparation (1 mM) was mixed with various concentrations of Acr, incubated for 1 h, and subjected to LC-MS/MS analysis. A, Extracted ion chromatograms of the mono-Acr adduct. B, Extracted ion chromatograms of the di-Acr adduct. C, Extracted ion chromatograms of the dehydrated products from the di-Acr adduct. The MS/MS spectrum of the corresponding parent ion is shown at the bottom of panels A–C.
